# DARPP-32 Distinguishes a Subset of Adult-Born Neurons in Zebra Finch HVC

**DOI:** 10.1101/2020.08.31.271783

**Authors:** Jake V Aronowitz, John R Kirn, Carolyn L Pytte, Gloster B Aaron

## Abstract

Adult male zebra finches (*Taeniopygia guttata*) continually incorporate adult-born neurons into HVC, a telencephalic brain region necessary for the production of learned song. These neurons express immediate early genes during song production, suggesting a role for neurogenesis in song production throughout the lifespan. Half of these adult-born HVC neurons (HVC NNs) can be backfilled from the robust nucleus of the arcopallium (RA) and are a part of the vocal motor pathway underlying learned song production, but the other half do not backfill from RA, and they remain to be characterized. Here we used cell birth-dating, retrograde tract tracing, and immunofluorescence to demonstrate that half of all HVC NNs express thephosphoprotein DARPP-32, a protein associated with dopamine (DA) receptor expression. We also demonstrate that DARPP-32+ HVC NNs are contacted by tyrosine hydroxylase immunoreactive fibers suggesting that they receive catecholaminergic input, have transiently larger nuclei than DARPP-32− HVC NNs, and do not backfill from RA. Taken together, these findings help characterize a group of HVC NNs that have no apparent projections to RA and so far have eluded any positive identification other than HVC NN status.

## Introduction

Songbirds, including zebra finches (*Taeniopygia guttata*), continually incorporate new neurons into brain regions that subserve song learning, perception, and production (Kirn & Devoogd, 1989; Sohrabji, et al., 1993; Alvarez-Buylla et al., 1988). The telencephalic nucleus HVC (proper name), which is necessary for learned song production, incorporates new neurons throughout adulthood, and these neurons express immediate early genes during song production (Jarvis, et al., 1998; Tokarev, et al., 2016). Approximately half of all new neurons that are added to HVC (HVC NNs) in adult male zebra finches send their axons to the robust nucleus of the arcopallium (RA), a premotor relay and the first synaptic target of the vocal motor pathway underlying song production (Figure 1A; Kirn, et al., 1991, 1999). In male zebra finches, 42% of HVC NNs can be backfilled from RA (HVC_RA_) at 30 days post-mitosis, and this increases to only ~50% at survival times from 3 months to 4 years after cell birth-dating (Scotto-Lomassese, et al., 2007; Walton, et al., 2012). This suggests that that there are two populations of new neurons added to HVC: those that project to RA and those that do not (Tokarev et al., 2016).

**Figure 1.**
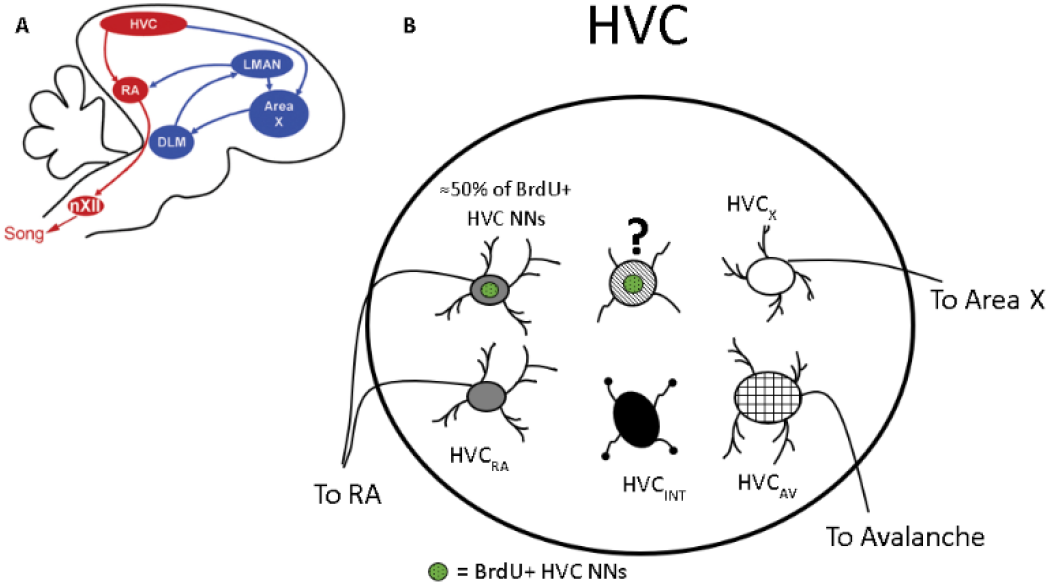
Organization of HVC and the zebra finch song system. (A) Schematic of a sagittal view of the zebra finch brain showing the song system brain regions that comprise the vocal motor pathway (red) and the anterior forebrain pathway (blue). Posterior is to the left, ventral is down. Reproduced with permission from Wood et al 2013. (B) Schematic of HVC showing known populations of HVC neurons. Green stippled circles are BrdU+ nuclei meant to distinguish those generated during our BrdU injections from those generated on days we did not inject BrdU, and therefore are BrdU-. The question mark represents the approximately 50% of HVC NNs that cannot be backfilled from RA.

Interestingly, the full repertoire of HVC neurons is not yet established. Four categories of HVC neurons have been identified based on projections and neurotransmitter markers: those that project to the striatal region Area X (HVC_X_), those that project to RA (HVC_RA_), a small population that project to the nucleus avalanche (HVC_AV_), and inhibitory interneurons (HVC_INT_; Dutar, et al., 1998; Mooney & Prather, 2005; Daou, et al., 2013; Ross, et al., 2017, 2019; Akutagawa & Konishi, 2010; Roberts, et al., 2017). Four morphological categories based on spine density and dendritic branching have also been described (Nixdorf et al.,1989) as well as four distinct electrophysiological classes (Kubota and Taniguchi, 1998). More recently, gene expression profiles revealed 5 distinct glutamatergic neuron types in HVC, suggesting 2 subcategories of the 3 projection neuron types (and 8 GABAergic subtypes; Colquitt et al., 2021). This echoes the proposition of two groups of putative “HVC_RA_^”^ neurons, one of which was based on axonal myelination (Egger, et al. 2020), and the other based on whether the cells responded to depolarizing current injection with tonic or phasic firing (Shea, et al., 2010).

This non-backfilling population of HVC NNs has long evaded further classification, largely because it has been assumed that HVC_RA_ neurons are the only class of adult-born neuron that is incorporated into HVC. HVC NNs do not project to Area X (Figure 1B; Alvarez-Buylla, et al., 1988). HVC_INT_ may be generated in HVC of juvenile male zebra finches (Scott & Lois, 2007), but this has not been replicated in adults (Scotto-Lomassese, et al., 2007; Walton, et al., 2012). Therefore, two broad explanations have been posited for the identity of HVC NNs that do not backfill from RA: 1) they are HVC_RA_ neurons that do not transport retrograde tracers to their soma for unknown reasons, or 2) they are not HVC_RA_ neurons, but rather project locally within HVC, or to some other brain region (e.g. Nucleus Avalanche, Av; Akutagawa & Konishi, 2010; Tokarev, et al., 2016). This suggests that HVC NNs that cannot be backfilled from RA are 1) excitatory interneurons, 2) local inhibitory interneurons that do not express any of the markers of inhibitory interneurons used in other studies (Wild, et al., 2005; Scotto-Lomassese, et al., 2007; Walton, et al., 2012), 3) Avalanche-projecting neurons (HVC_AV_), or 4) HVC_RA_ neurons that fail to transport the retrograde tracer, perhaps due to their axon or synapse type, or due to technical limitations of the tracer. Tokarev and colleagues showed that HVC NNs that were not retrogradely labeled from RA were found 3-, 5-, and 8-weeks post cell birth-dating (Tokarev, et al 2016). Intriguingly, they also showed that these non-backfilled HVC NNs expressed IEGs after song production, and on this basis they postulated that there may a second type of HVC NN that demonstrates song-related activation but may never synapse onto RA.

Here, we used cell birth-dating, immunofluorescence, and retrograde tracing to characterize the HVC NNs that do not backfill from RA. First, we show that nearly half of HVC NNs express DARPP-32 (“DARPP-32+ HVC NNs”). DARPP-32 (DA and cAMP regulated phosphoprotein, 32 kDa) is a phosphoprotein located in the nucleus, cytoplasm, axons, axon terminals, and dendrites of dopaminoceptive neurons (Walaas & Greengard, 1984; Ouimet & Greengard, 1990; Svenningson, et al., 2004; Stipanovich, et al., 2008). We also demonstrate that DARPP-32+ HVC NNs do not backfill from RA at 32 days post-mitosis, have transiently larger nuclei than DARPP-32− HVC NNs, and are contacted by tyrosine hydroxylase-immunoreactive (TH+) fibers. We propose that DARPP-32+ HVC NNs comprise a unique subpopulation of HVC NNs and represent the 50% of HVC NNs in adult-male zebra finches that historically could not be retrogradely labeled from RA.

## Materials and Methods

### Animals

All procedures were performed in accordance with the Wesleyan University Institutional Animal Care and Use Committee (IACUC) regulations. Adult male zebra finches (0.5 to 1.5 years old) were hatched in the Wesleyan University breeding colony. Females were not included in this study because the zebra finch song system is sexually dimorphic, and female HVC is small and does not receive many new neurons in juveniles or adulthood (Burek, et al., 1994; DeWulf & Bottjer, 2005; Shaughnessy, et al., 2019). Birds in this study came from individual family breeding cages with one nest, or from larger cages with two nests. Birds reared in either of these conditions were balanced across groups to control for any effect of juvenile housing condition. When birds reached 90 days post hatch (dph), they were moved from the breeding colony into larger cages where they were group-housed with similarly aged males. Songbirds housed with other males show greater proliferative activity (Shevchouk, et al., 2017), and so we housed our experimental birds in auditory and visual contact of other males to maximize new neuron production. Zebra finches used for time course dependent DARPP-32 expression experiments (n=43, 100-555 dph) were randomly distributed into 8 groups that differed in the amount of time between final BrdU injection and the date of sacrifice. Tissue from birds in this first cohort was also used for tyrosine hydroxylase experiments (n=3, age range: 169-203 dph) and nuclear diameter experiments (n=10, age range: 164-301 dph). For retrograde tracing experiments, we used a separate group of adult male zebra finches (n=4, age range: 144-175 dph).

### BrdU Injections

Zebra finches received intramuscular (I.M.) injections of 2-Bromo-5’-deoxyuridine (BrdU; Roche Diagnostics) in 0.05 M tris buffered saline (TBS; pH=7.4; 15 mg/ml, approx. dosage 46.9 – 57.7 mg/kg) into the pectoralis muscle twice daily for three consecutive days to label dividing cells. BrdU is bioavailable for approximately 1-2 hours following I.M. injection (Barker, et al., 2013). We performed the first injection ~ 10 AM and the second injection ~ 5 PM to avoid differences in circadian rhythms that could affect the amount of new neuron labeling.

### Stereotaxic Surgery

28 days following the last BrdU injection, birds (n=4, Fig. 5) received I.M. injections of meloxicam (2 mg/kg) followed by anesthesia via I.M. injections of ketamine and xylazine (0.03 ml of 10 mg/ml ketamine; 0.03 ml of 20 mg/ml xylazine). They were then placed in a customized stereotaxic apparatus on a heating pad with their head at a 45◻ angle to the horizontal plane. A midline incision was made on the dorsal surface of the head to expose the skull. A small craniotomy (~0.5 mm^2^) was then made above the bifurcation of the midsagittal sinus (Y-point) and the stereotaxis was ‘zeroed’ at this Y-point (Nixdorf-Bergweiler & Bischof, 2007). Small craniotomies (~0.5 mm^2^) were made over the injection sites for RA: A/P = −1.7 mm and −1.9 mm; M/L= ± 2.65 mm; D/V= −2.3 mm and −2.5 mm, for a total of 4 injection sites per hemisphere. Injections were made at a 9◻ angle posteriorly off the vertical axis to avoid damage to HVC en route to RA. The pia was then gently punctured with a 26.5-gauge beveled syringe needle. A 2% Fluorogold solution in 0.9% NaCl (“FG”; Fluorochrome, Inc.) was slowly pressure injected, ~20 nl per site, four sites per RA. Following injections, the craniotomies were filled with Kwik-Cast (World Precision Instruments, Inc.), the incision was sutured, antibacterial/analgesic ointment applied, and the animal placed in a surgical recovery cage under a heat lamp with food and water available *ad libitum* until the effects of anesthesia had subsided, at which time they were returned to their presurgical cages.

### Histology/Immunofluorescence

At varying survival times following the last BrdU injection (days post injection, dpi), birds were deeply anesthetized with a mixture of ketamine (0.07 ml of 10 mg/ml solution; Ketaset) and xylazine (0.07 ml of a 20 mg/ml solution; Xylamed). When birds were unresponsive to a toe-pinch they were transcardially perfused with 20 ml 0.1M PBS (pH=7.4) followed by 30 ml 4% paraformaldehyde (PFA, pH=7.4; Electron Microscopy Sciences). Brains were carefully dissected, post-fixed for 24 hrs. in 4% PFA at 4◻C, and sectioned sagittally at 50 μm on a vibratome (Leica VT1000S). We found that DARPP-32 labeling was most robust and antibody binding the most specific when sections were processed within 1-2 weeks after perfusion without cryoprotection for long term storage. Starting with the first complete section through HVC, every 3^rd^ section containing HVC was used for IHC analysis. Free-floating sections were permeabilized for 30 min in 0.5% PBS-T (PBS with Triton-X 100) at room temperature on a shaker. Then, antigen retrieval was performed via 20 min incubation in 1.5 N HCL at 37◻C in a water bath, followed by neutralization with two, 5 min rinses in Tris base (pH=8.56; Fisher Scientific). Sections were then blocked with normal goat serum and normal donkey serum in PBS-T (3% each) for 20 min at room temperature on the shaker. Sections were incubated overnight at 4◻C in varying mixtures of the following primary antibodies: rabbit anti-DARPP-32 (1:1000; Abcam Cat# ab40801, RRID:AB_731843), mouse anti-Hu (1:500; Molecular Probes, Invitrogen Cat# A-21271, RRID:AB_221448), rat anti-BrdU (1:500; Bio-Rad Cat# MCA2060, RRID:AB_323427), and mouse anti-tyrosine hydroxylase (1:2000; Millipore Cat# MAB5280, RRID:AB_2201526). Following rinses in PBS-T, sections were incubated at room temperature for 90 min in a cocktail of the following secondary antibodies: goat anti-rabbit Alexa Fluor 555 (1:500; Molecular Probes, Invitrogen, Cat# A-21429, RRID:AB_2535850), donkey anti-mouse Alexa Fluor 647 (1:500; Molecular Probes, Invitrogen, Cat# A-31571, RRID:AB_162542), and goat anti-rat Alexa Fluor 488 (1:500; Molecular Probes (Invitrogen) Cat# A-11006, RRID:AB_141373). Following 3 PBS-T rinses, sections were counterstained with Hoechst 33258 (2 ug/ml in PBS; Molecular Probes), except for tissue from the HVC_RA_ backfill experiments, which was not counterstained because FG fluoresces in the same channel as Hoechst 33258 and the overlap would have hindered interpretation. Finally, the sections were rinsed in PBS, mounted on positively charged slides (Superfrost++, Fisher Scientific), and coverslipped with Aqua-Poly/Mount (Polysciences, Inc.). Slides were then photoprotected and stored at −20◻C until confocal imaging. Negative controls were performed in which one or more primary antibodies were omitted to check for cross-reactivity.

### HVC Analyses

HVC was traced using computer-yoked microscopy software (Neurolucida; MicroBrightField, Inc.), as defined by DARPP-32 expression, which was much more robust around HVC and sparser within it (Figure 2; Singh & Iyengar, 2019). These boundaries corresponded to the previously defined cytoarchitectural boundaries of HVC described by Fortune & Margoliash (1995), were also identifiable with dark field microscopy, and with fluorescent microscopy to visualize the neuron-specific protein Hu. All images were analyzed in Leica Application Suite-X software (LAS− X; Leica, Inc.). Nuclear diameters were measured in LAS-X at the widest portion of the nucleus. All experimenters were blind to animal ID, hemisphere, and condition to ensure a lack of bias and maximum reproducibility. As mentioned above, we sampled every third 50 μm section (every 150 micrometers) throughout the mediolateral extent of HVC which eliminates the possibility of double counting cells that may have been split in half between adjacent sections. In each section, we traced the boundaries of HVC (as defined by DARPP-32 expression) and counted all DARPP-32+ DARPP-32− HVC NNs within those boundaries. Cells were included in our counts if their nucleus contacted the traced boundary of HVC. A 3-μm-thick buffer layer was applied to the top and bottom of every Z-stack (on the Z-axis). Cells were not counted if the broadest cross section of their nucleus was in focus in either buffer layer. To calculate new neurons per cubic millimeter, we measured the area of HVC in each slice and multiplied that value by the section thickness and sampling interval. DARPP-32 undergoes constant cytonuclear shuttling as a function of its phosphorylation state (Stipanovich, et al., 2008). Our antibody against DARPP-32 binds to a 16 AA segment on a region of the protein that does not contain any phosphorylation sites (Abcam ab40801) and therefore it labels DARPP-32 irrespective of phosphorylation state and cellular location. Consistent with this, in HVC we observed DARPP-32 labeling of 1) somata + nuclei, 2) somata only, 3) nuclei only. Due to this, we decided to count all DARPP-32+ HVC NNs regardless of the cellular location of DARPP-32.

**Figure 2.**
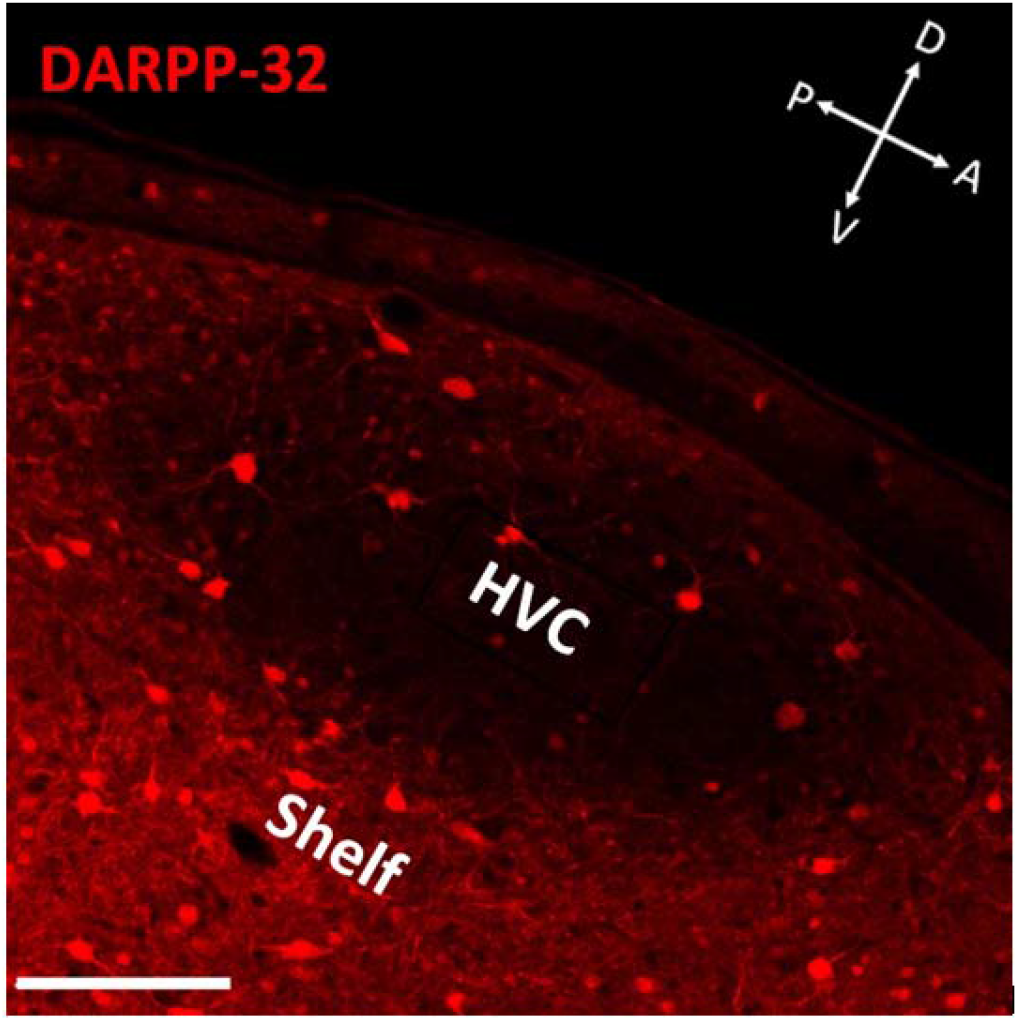
Confocal photomicrograph of HVC in sagittal section showing DARPP-32 labeling using a monoclonal antibody (ab40801) that recognizes and binds to DARPP-32 independent of its current phosphorylation state. There are fewer labeled cells in HVC compared to the underlying HVC shelf region, which, like much of the surrounding nidopallium, labels much more robustly for DARPP-32 than within HVC (Singh & Iyengar, 2019). 23.8X, scale bar = 100 μm.

### Confocal Microscopy

Fluorescent images were captured using a Leica DMI8 inverted confocal microscope (Leica Microsystems) Every third section spanning the mediolateral extent of HVC was used for IHC and subsequently imaged with a 5x, 10x, 20x, 40x, or 63x objective using excitation lasers at 405 nm, 488 nm, 552 nm, and 635 nm. Confocal imaging parameters were saved as a user file and these parameters were kept consistent for all slides imaged within a particular experiment.

### Experimental Design and Statistical Analysis

We used 8 groups varying in survival time after the final BrdU injection (7 – 403 dpi) in order to determine whether DARPP-32 expression differed across new neuron age. The 7, 14, 21, 90, and 365-403 dpi groups had n=5, the 180 dpi group had n=4, the 28 dpi group had n=6, and the 45 dpi group n=7. We did not see differences between 365 and 403 dpi animals, so we combined these animals into one group. We report the results of a one-way ANOVA for 8 samples, and Bonferroni post hoc tests to adjust for multiple comparisons, both performed in SPSS statistical software (IBM). We ran one-way ANOVAs to compare 1) the *fraction* of all HVC NNs that expressed DARPP-32, calculated as DARPP-32+ HVC NNs / total HVC NNs (Figure 4B), and 2) the *density* of DARPP-32+ HVC NNs per cubic millimeter (Figure 4A). For retrograde tracing studies evaluating HVC NN connectivity, we first turned off DARPP-32 and Hu fluorescent channels in LASX and then a blinded experimenter marked all BrdU+/FG− and BrdU+/FG+ cells before turning on the other two channels to determine whether they also expressed DARPP-32, Hu, or both. For both DARPP-32+ and DARPP-32− HVC NNs, we calculated the fraction of the total that possessed FG, i.e. projected to RA (371 cells from 7 hemispheres from 4 birds). For nuclear diameter experiments (Figure 7), we first ran a two factor ANOVA with repeated measures on cell type. Then, we ran paired T-tests to compare the difference between neuron types at a given dpi, and we used one-way ANOVAs with Tukey’s post hoc tests to analyze changes in the nuclear diameter of HVC NNs as a function of days post BrdU injection (709 cells from 20 hemispheres from 10 birds). These were performed in SPSS.

**Figure 3.**
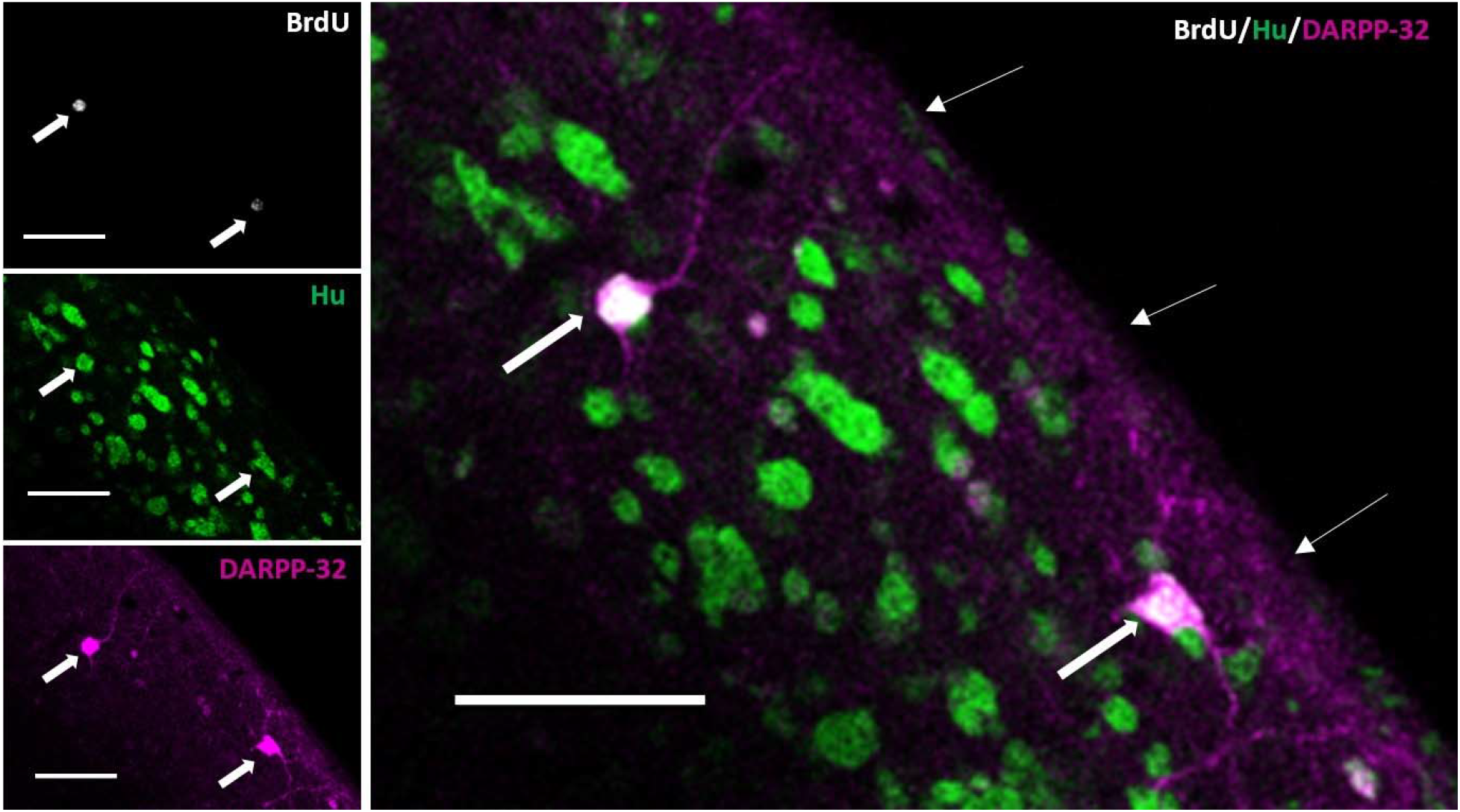
28-day old HVC NNs express the phosphoprotein DARPP-32. Confocal photomicrograph of a single optical plane of section of the same field of view in dorsal HVC showing two DARPP-32+ NNs at 28 dpi. **(A)** BrdU labeled nuclei. **(B)**Neurons labeled for the neuron-specific protein Hu. **(C)** DARPP-32+ neurons. **(D)** merged image of all three channels; thick arrows indicate triple-labeled BrdU+/Hu+/DARPP-32+ HVC NNs at 28 dpi. Thin arrows indicate the dorsal surface of HVC. 18x magnification. Scale bars = 25 μm.

**Figure 4.**
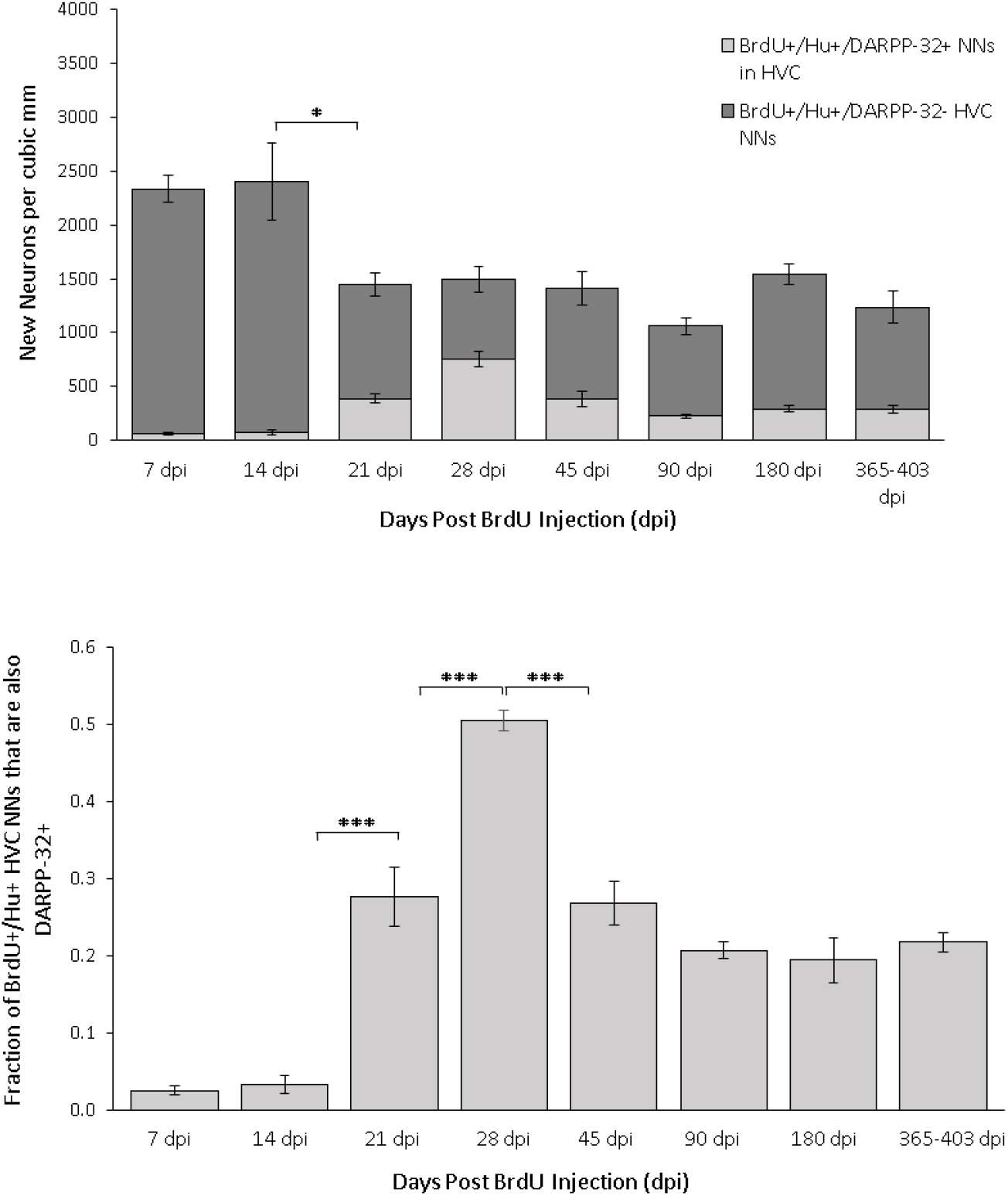
DARPP-32 expression in HVC NNs changes as a function of neuronal age. **(A)** The density of HVC NNs as a function of days following the final BrdU injection. Light gray bars represent BrdU+/Hu+/DARPP-32+ HVC NNs, dark gray bars represent BrdU+/Hu+/DARPP 32− HVC NNs. The total height of the stacked bars equals the average density of HVC NNs per cubic millimeter. We observed a significant decline in the density of HVC NNs between 14 and 21 dpi (t(8) = 2.57, p = 0.03, independent samples T-test). After 21 dpi we observed no group differences in the density of HVC NNs. Error bars represent SEMs. Group sizes are as follows: 7, 14, 21, 90, and 365 dpi, n=5; 28 dpi, n=6; 45 dpi, n=7; 180 dpi, n = 4. **(B)** To determine the fraction of HVC NNs that express DARPP-32 at each time point sampled, we divided the density of DARPP-32+ HVC NNs (BrdU+/Hu+/DARPP-32+) by the density of all HVC NNs (BrdU+/Hu+); these values are presented here as a derivation of 4A. These data show that the fraction of HVC NNs that express DARPP-32 increases to its peak at 28 dpi (50.4 ± 1.6%), then declines to 26.8 ± 7.3% at 45 dpi. At all later time points sampled, the fraction of HVC NNs expressing DARPP-32 was not significantly different from the 45 dpi time point, indicating that the number of DARPP-32+ HVC NNs in a given cohort of NNs is stable from 45 dpi onward. Error bars represent SEMs. Significance markers are as follows: p<.05 = * p<.01 = **; p<.001 = *** and these markers are consistent throughout the manuscript.

## Results

### HVC NNs May Transiently Express DARPP-32

We found that approximately half of all HVC NNs express DARPP-32 at 28 dpi (50.4 ± 3.2%; Figures 3, 4A, 4B). To see whether DARPP-32 expression in HVC NNs changes as a function of neuronal age, we examined HVC NNs at eight time points ranging from 7 to 403 days after the last BrdU injection. We found a significant effect of neuronal age on the fraction of HVC NNs that express DARPP-32 (F(7,34) = 46.9, p <.001; Figure 4B), and on the density of DARPP-32+ HVC NNs per mm^3^ (F(7,34) = 19.8, p <.001; Figure 4A). In the first month post-mitosis, we observed an increase in the fraction of HVC NNs that express DARPP-32 (Figure 4B). Tukey’s post-hoc tests performed on DARPP-32+ HVC NN fractions revealed significant differences between 7 and 21 dpi (p <.001), 14 and 21 dpi (p <.001), 21 and 28 dpi (p <.001), indicating that the number of DARPP-32+ HVC NNs in a given cohort of HVC NNs increases during the first month post mitosis. From 28 to 45 dpi we observed a decline in the fraction of DARPP-32+ HVC NNs (p <.001), after which there was no change in the fraction of HVC NNs that expressed DARPP-32 (Figure 4B). During the same time, over days 28-45 dpi, there was no decline in total number of BrdU birth-dated HVC NNs. This suggests that a subset of HVC NNs may transiently express DARPP-32 during their first month of development, and thereafter turn off DARPP-32 expression.

### DARPP-32+ HVC NNs Do Not Backfill from RA at 32 Days Post-Mitosis

To test whether DARPP-32+ HVC NNs project to RA, we birth-dated NNs with BrdU and injected FG into RA (Figure 5A-C). Birds were killed at 32 dpi, 4 days following surgery. Walton et al (2012) reported that 42% of HVC NNs project to RA at 30 dpi. Similarly, we found that 46.4% of HVC NNs examined were RA-projecting (Figure 5D-I). When we specifically examined RA-projecting HVC NNs, we observed that they were nearly exclusively DARPP-32− (171/172 cells, 99.4%; Figure 5I). Moreover, all but one (125/126) DARPP-32+ HVC NNs examined were FG−, demonstrating that DARPP-32+ HVC NNs are not backfilled from RA at 32 dpi and thus may not be RA-projecting.

**Figure 5.**
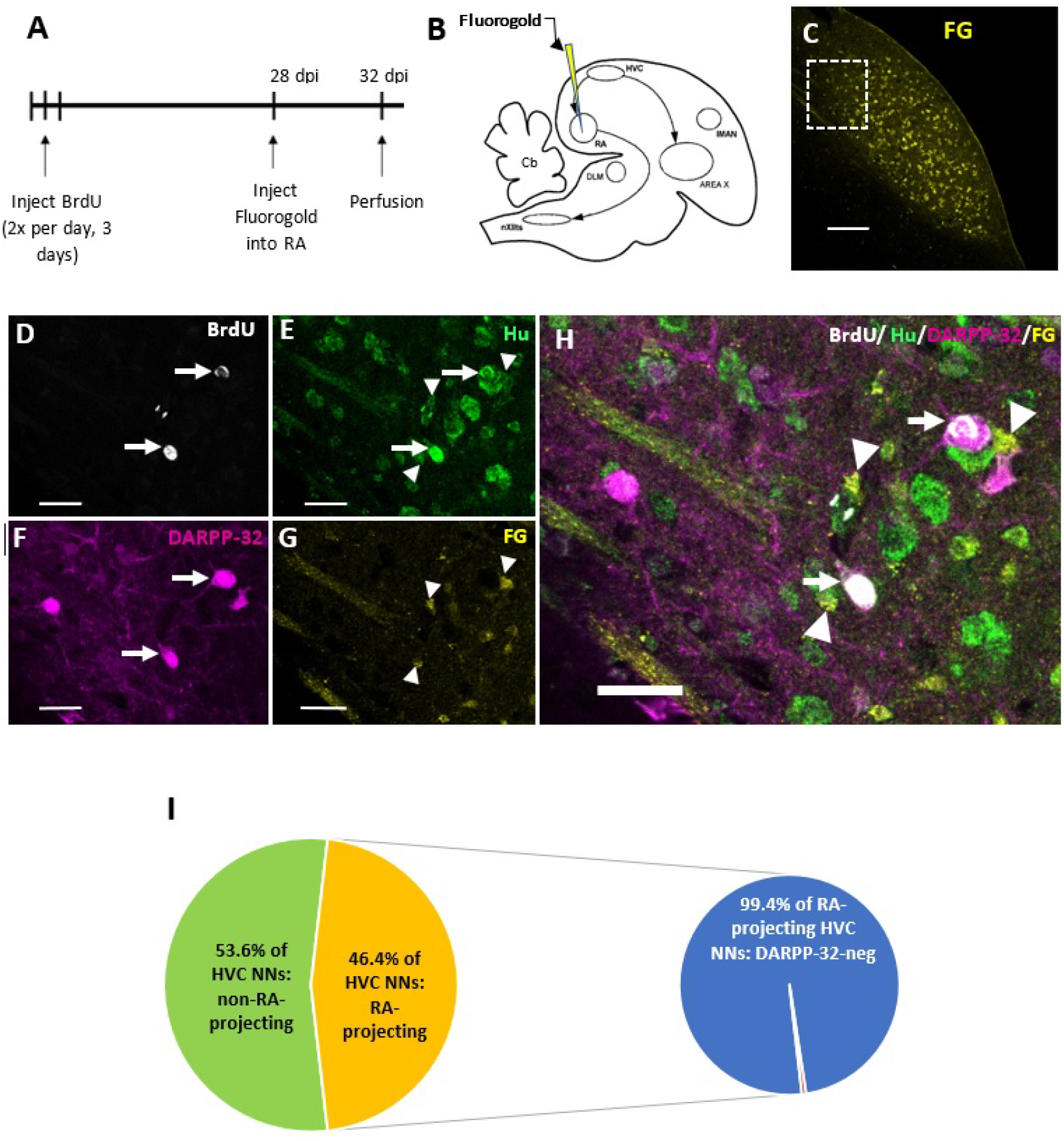
DARPP-32+ HVC NNs do not backfill from RA at 32 dpi. **(A)** Timeline of HVC_RA_ backfill experiments. **(B)** Simplified schematic of the song system in a sagittal section showing some of the brain regions involved in song production. The retrograde tracer Fluorogold (FG) was injected into RA to backfill HVC_RA_ neurons. **(C)** Maximum intensity projection of a Z-stack showing backfilled FG+, RA-projecting neurons in HVC (18x; scale bar = 100 μm). Dotted white box delineates the area in HVC examined at higher magnification in panels D-H (18x, scale bar = 100 μm). **(D-H)** Confocal photomicrograph showing DARPP-32+ HVC NNs (white arrows) in HVC that did not backfill from RA. **(D)** BrdU+ nuclei. **(E)** Hu+ neurons. **(F)** DARPP 32+ neurons. **(G)**FG+ neurons. **(H)**Merged image of all four channels showing DARPP-32+ HVC NNs that do not backfill from RA (white arrows), surrounded by Hu+/FG+/DARPP-32 HVCRA neurons (white arrowheads). 50x, scale bar = 25 μm. **(I)**125/126 DARPP-32+ HVC NNs we examined were FG-negative. 32-34 day old DARPP-32+ HVC NNs do not backfill from RA. 53.6% of HVC NNs examined (green; 199/371) did not project to RA, while 46.4% did project to RA (yellow; 172/371 neurons), as measured by whether or not the somata were FG+. Of the 46.4% of RA-projecting HVC NNs, only 1/172 was DARPP-32+, suggesting that DARPP-32+ HVC NNs do not backfill from RA at 32 dpi.

### DARPP-32+ HVC NNs Receive Catecholaminergic Input

DARPP-32 is a reliable marker of dopaminoceptive neurons in both mammals (Hemmings, et al., 1987) and zebra finches (Rochefort, et al., 2007; Kosubek-Langer, et al., 2017). To determine whether DARPP-32+ HVC NNs receive catecholaminergic input, we labeled tyrosine hydroxylase, the rate-limiting enzyme in the catecholamine synthetic pathway (Fernstrom & Fernstrom, 2007). At 28 dpi, all DARPP-32+ HVC NNs we examined appeared to be contacted by TH+ fibers (Figure 6; 74 cells from 3 birds). These data strongly suggest that DARPP-32+ HVC NNs receive dopaminergic input, likely from the periaqueductal gray (PAG), which is the primary source of dopaminergic afferents to HVC (Appeltants, et al., 2001; Tanaka, et al., 2018).

**Figure 6.**
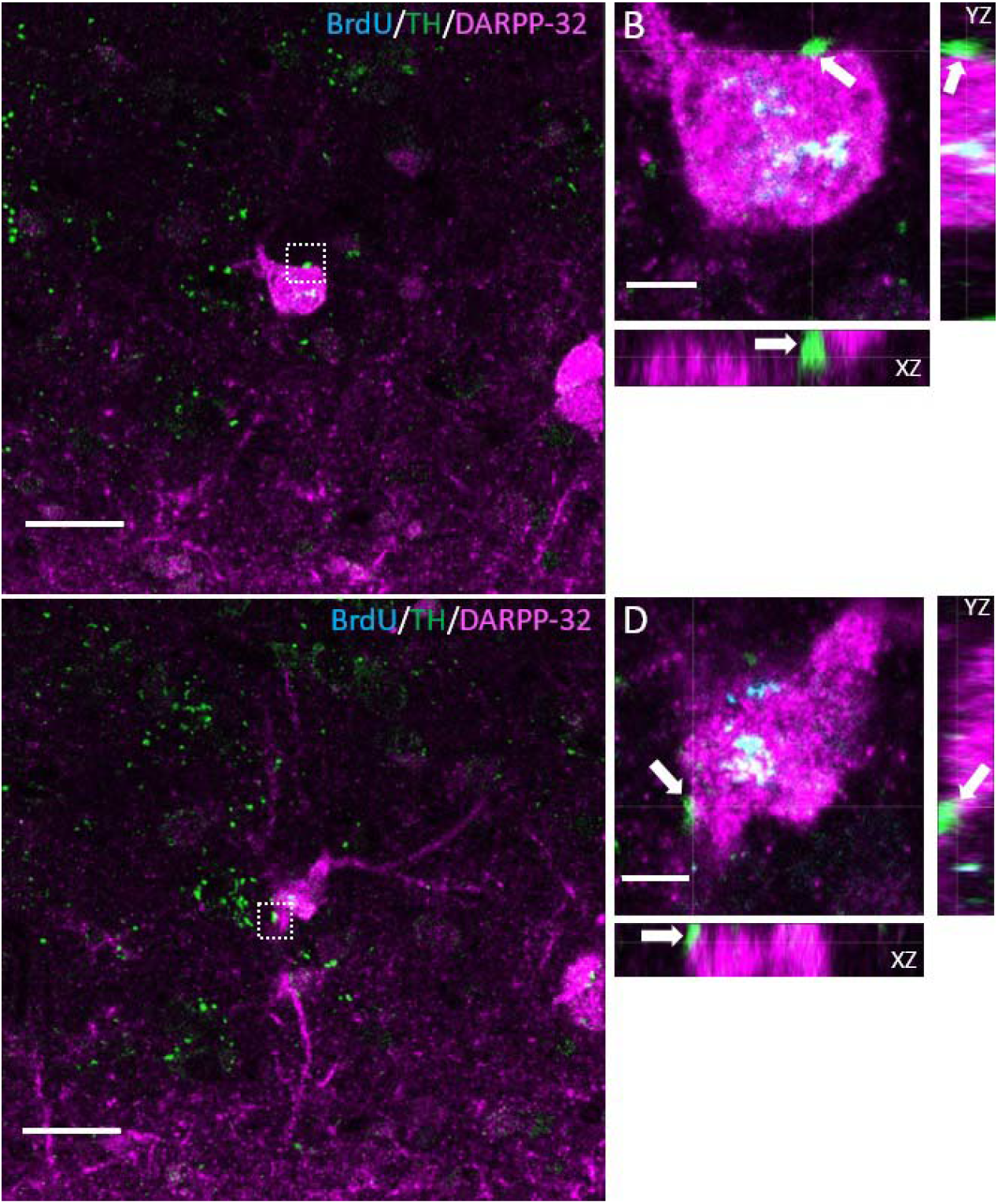
DARPP-32+ HVC NNs are contacted by tyrosine hydroxylase fibers. **(A,C)** Confocal photomicrographs of the same DARPP-32+ HVC NN at two different levels in the Z-stack, showing two different putative TH synaptic contacts. **(B,D)** High magnification orthogonal views of the synaptic contacts shown in the dotted white boxes in images **A**and **C**. Scale bars: 10 μm: **(B, D)**, 25 μm **(A, C)**. Magnification: 75.2x **(A,C)**, 140x (**B,D)**.

### DARPP-32+ HVC NN Nuclear Diameters are Transiently Larger than DARPP-32− HVC NN Nuclear Diameters

We noticed qualitatively that DARPP-32+ HVC NN nuclei appeared to be larger than those of DARPP-32− HVC NNs. To test this quantitatively, we measured the nuclear diameter of DARPP-32+ and DARPP-32− HVC NNs at 21, 28, and 45 dpi. Previous work in canaries has shown that the nuclear diameter of HVC NNs – whether retrogradely labeled from RA or not – decreases between 30 and 240 dpi (Kirn, et al., 1991). In a subset of our birds (n=709 neurons from 20 hemispheres from 10 birds) we evaluated whether the nuclear diameter of DARPP-32+ HVC NNs changed as a function of neuronal age, and whether it differed from DARPP-32− NNs (Figure 7). Two factor ANOVA with repeated measures on cell type revealed a significant main effect of neuronal age (F(9) = 22.04, p < 0.001), a significant main effect of cell type (F(10) = 227.4, p <.001), and a significant interaction between neuronal age and cell type (F(19) = 15.9, p <.01). We next evaluated how the nuclear diameter of each cell type changed with time. One way ANOVAs revealed that the nuclear diameter of both DARPP-32− HVC NNs (F(2,7)=10.8, p <.01) and DARPP-32+ HVC NNs (F(2,7)=27.9, p <.001) decreased significantly as the neurons matured (Figure 7). Tukey’s post-hoc tests showed that the nuclear diameter of DARPP-32− HVC NNs decreased significantly between 28 and 45 dpi (p <.01), while the nuclear diameter of DARPP-32+ HVC NNs decreased between 21 and 45 (p <.01), as well as 28 and 45 dpi (p <.01). Finally, we showed that DARPP-32+ HVC NNs had significantly larger average nuclear diameter than DARPP-32− HVC NNs at 21 dpi (t(4) = 4.53, p <.05) and at 28 dpi (t(4) = 11.31, p <.001). At 45 dpi, however, there was no difference between the average nuclear diameters of the two putative subtypes of HVC NNs (t(4) = 1.97, p >.05).

**Figure 7.**
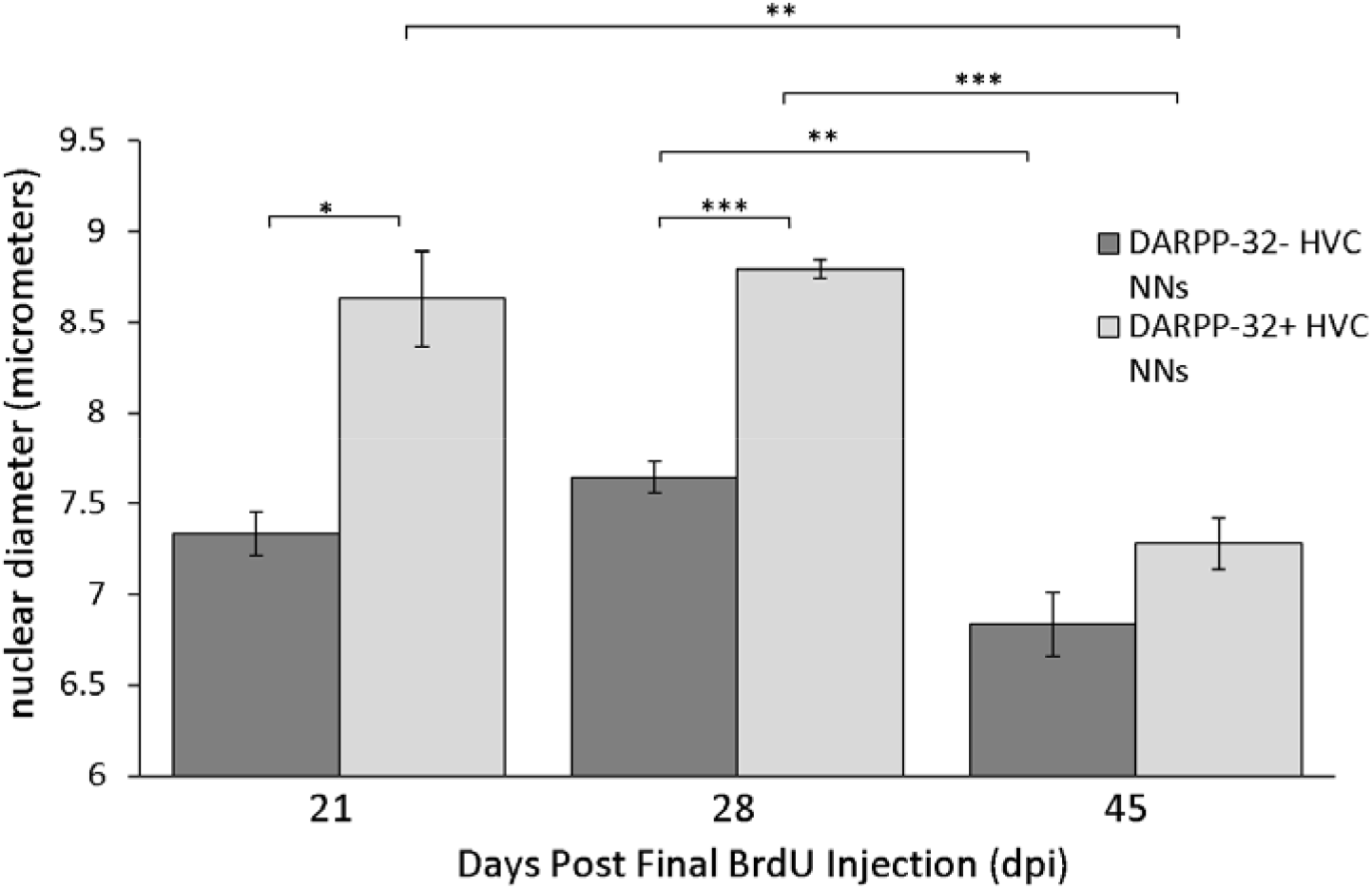
Mean nuclear diameter of HVC NNs over the first 45 days post-BrdU injection. The mean nuclear diameter of DARPP-32+ HVC NNs (light gray bars) was significantly larger than that of DARPP-32− HVC NNs (dark gray bars) at 21 dpi, and at 28 dpi. However, at 45 dpi, there was no significant difference between the mean nuclear diameter of the two subtypes of HVC NNs. Statistics revealed that both subtypes of new neurons demonstrated a decline in nuclear diameter with increasing neuronal age.

## Discussion

We characterize a subtype of HVC NN that expresses DARPP-32 and does not backfill from RA. We demonstrate that DARPP-32+ HVC NNs are putatively contacted by TH+ fibers and have transiently larger nuclei than DARPP-32− HVC NNs. In both canaries (Alvarez-Buylla, et al., 1988) and zebra finches (Walton, et al., 2012; Tokarev, et al., 2016) there is a fraction of HVC NNs that neither backfill from RA, nor express common markers of HVC inhibitory interneurons (Scotto-Lomassese, et al., 2007). Since Area X projecting HVC neurons are only generated prenatally (Alvarez-Buylla, et al., 1988; Scharff, et al., 2000), this population was commonly explained as young neurons not yet transporting the tracer (Kirn, et al., 1991, 1999; Scotto-Lomassese, et al., 2007). However, this is unlikely because nearly 4 years post-BrdU injection, many HVC NNs still did not backfill from RA, suggesting that they may never develop connections to RA or they never transport the tracer (Walton, et al., 2012).

Tokarev and colleagues (2016) showed that HVC NNs that do not backfill from RA express immediate early genes following song production at 3-, 5-, and 8-weeks post BrdU injections (Tokarev, et al., 2016). From this, they postulated that there is a second type of neuron added to adult HVC that participates in song production and may not project to RA (Tokarev, et al., 2016). In support of those data, we show that one-month old DARPP-32+ HVC NNs do not backfill from RA. We propose that DARPP-32+ HVC NNs are either 1) excitatory HVC interneurons, 2) inhibitory interneurons that do not express the calcium-binding proteins that have traditionally been used to tag HVC_INT_ (parvalbumin (PV), calretinin (CR), and calbindin (CB); Wild, et al., 2005; Scotto-Lomassese, et al., 2007), 3) HVC_AV_ projection neurons (HVC neurons that send their axons to nucleus Avalanche), or 4) HVC_RA_ neurons that fail to transport the tracer due to their synapse or axon type. Option 2 is unlikely because the number of GABA+ HVC_INT_ reported in HVC is not significantly different from the number of HVC_INT_ labeled with PV, CR, and CB combined (Wild, et al., 2005; Scotto-Lomassese, et al., 2007). Option 3 is unlikely because HVC_AV_ neurons are too sparse to account for the ~50% of HVC NNs that do not backfill from RA. Roberts et al (2017) reported that HVC_AV_ neurons comprise < 1% of all HVC neurons. We estimate the total number of BrdU+/Hu+/DARPP-32+ NNs added to HVC per hemisphere during three days of BrdU injections to be 189.2 ± 17.4, based on our 28 dpi values and assuming an HVC volume of 0.25 mm^3^, following Walton, et al., (2012). Roberts et al., (2017) estimated the total number of HVC_AV_ neurons to be 130 ± 69.8 per hemisphere. Though we cannot presently rule it out, option 4 is unlikely because Fluorogold is a highly effective retrograde tracer that rapidly (< 1 week) labels neurons as efficiently, or more efficiently (Yao, et al., 2015), than other types of retrograde tracers (e.g. Cholera toxin subunit B, Fast Blue, Fluoro-Ruby, DiI, etc). However, we know of no published studies that show evidence of a retrograde tracer labeling every member of a projection neuron population. While it is possible that DARPP-32+ HVC NNs simply failed to transport the tracer due to their synapse or axon type, we believe the more likely explanation is that DARPP-32+ HVC NNs are not RA-projecting but in fact project locally as excitatory interneurons. In support of this idea, a recent transcriptomic study of the vocal motor pathway in zebra finches revealed the presence of a neuron at the end of the neurogenic lineage (i.e. close to maturity) that expresses PPP1R1B (the gene encoding DARPP-32), NECTIN3 (gene encoding a cell adhesion molecule related to synapse formation and maintenance), and also SLC17A6 (a gene encoding the vesicular glutamate transporter, used as a marker of mature glutamatergic neurons; Colquitt, et al., 2021). This characterization fits nicely with the interpretation that DARPP-32+ HVC NNs are excitatory interneurons.

In Area X, a brain region necessary for song learning and maintenance, there is a steady increase in the proportion of adult-born neurons that express the immediate early gene EGR-1 (ZENK) and DARPP-32 between 21 and 42 dpi, suggesting a link between new neuron activation, DARPP-32 expression, and successful maturation of adult-born Area X neurons (Keilani, et al., 2012; Kosubek-Langer, et al., 2017). We found that between 7 and 28 dpi the fraction of HVC NNs expressing DARPP-32 increased from ~2.5% to 50% and then decreased until 45 dpi, after which time there was no further change in the fraction of HVC NNs that were DARPP-32+ (Figure 4B). Despite a decline in the fraction of HVC NNs expressing DARPP-32 between 28 and 45 dpi, there was no corresponding decline in the overall number of new neurons in HVC (Figure 4A). Taken together these data suggest that DARPP-32+ HVC NNs are not disproportionately dying. Rather it suggests that a subset of HVC NNs is shutting off DARPP-32 expression sometime after 28 dpi. Furthermore, it appears that while some new neurons that express DARPP-32 during maturation turn it off, others maintain expression for up to one year post mitosis.

To determine whether DARPP-32+ HVC NNs receive catecholaminergic inputs, we combined BrdU, DARPP-32, and TH immunofluorescence. At 28 dpi, all DARPP-32+ HVC NNs we examined were contacted by TH+ fibers, indicating that they receive catecholaminergic input by one-month post-mitosis. HVC receives dopaminergic inputs predominantly from the PAG and to a lesser extent, the ventral tegmental area (VTA, Appeltants, et al., 2000, 2001; Tanaka, et al., 2018). In addition to multiple types of DA receptors (Kubikova, et al., 2009), HVC contains α2-noradrenergic receptors and receives noradrenergic input from the midbrain (Riters, et al., 2002; Appeltants, et al., 2000). Hara et al. (2010) showed that α2- and β1-noradrenergic receptor activation can alter the phosphorylation state of DARPP-32 at threonine-34 (thr-34) in mouse neostriatal brain slices, suggesting that dopamine and norepinephrine (NE) both play a role in modulating DARPP-32 phosphorylation (Hara, et al., 2010). Thr-34 is also the residue at which DARPP-32 is phosphorylated by protein kinase A following DA receptor activation (Svenningsson, et al., 2004). This raises the possibility that DARPP-32+ HVC NNs are modulated by dopaminergic and/or noradrenergic inputs. However, our TH immunofluorescence procedures did not allow us to distinguish between dopaminergic and noradrenergic terminals.

While it is impossible to describe the function of these DARPP-32+ HVC NNs at this time, we offer some speculation that may help guide future discussions and studies. With regard to the role of DA in HVC: in juvenile zebra finches, DA from the PAG released onto HVC signals the social salience of the tutor and facilitates song copying (Tanaka, et al., 2018). In adult male canaries, pharmacological inactivation of the PAG significantly increases the latency to sing without affecting other aspects of song, suggesting that PAG_HVC_ dopamine is related to the motivation to sing in adult songbirds (Haakenson, et al., 2020). We suggest that DARPP-32+ HVC NNs receive DA from PAG and serve in the pathway coding for social salience as it relates to song production. DARPP-32+ HVC NNs do not backfill from RA at 32 days post-mitosis and therefore may not project to RA as part of the song motor pathway. Thus, while DARPP-32+ HVC NNs may be activated by PAG inputs related to the motivation to sing, they would not be able to affect a motor output directly, implying a relatively intermediary role in song production with respect to HVC_RA_ NNs. We also consider that the putative 50% split of HVC NNs into HVC_RA_ versus DARPP-32+ categories may reflect a functional one-to-one relationship between these groups. Considering all these implications together, we propose that DARPP-32+ NNs are added to HVC to serve as dopaminoceptive chaperones for HVC_RA_ NNs, receiving and relaying the social salience signals sent from the PAG.

In addition to signaling an increased motivation to sing, the presence of a female produces song that has more introductory notes, greater number of song motifs per bout, and is sung faster than undirected song (Sossinka & Bohner, 1980). Evidence suggests that HVC is responsible for the temporal patterning of song (Long and Fee, 2008). It is possible that activation of DARPP-32+ HVC NNs, and perhaps other DA receptor containing cells in HVC, serves to alter HVC network properties in a way that produces a faster version of the song with more introductory notes and more motifs per bout (Sossinka & Bohner, 1980). As suggested above, we propose that DARPP-32+ HVC NNs may serve at least partly as intermediaries between the DA signal received from PAG and their HVC_RA_ NN partners.

## Acknowledgements

The authors would like to thank Dr. Luke Remage-Healey for his helpful comments on earlier versions of the manuscript.

## References

Alvarez-Buylla A, Theelen M, & Nottebohm F (1988). Birth of projection neurons in the higher vocal center of the canary forebrain before, during, and after song learning. Proc. Natl. Acad. Sci. USA, 85:8722–8726.

Alvarez-Buylla A, Kirn JR, Nottebohm F (1990) Birth of projection neurons in adult avian brain may be related to perceptual or motor learning. Science 249(4975):1444–1446.

Appeltants D, Absil P, Balthazart J, Ball GF (2000) Identification of the origin of catecholaminergic inputs to HVc in canaries by retrograde tract tracing combined with tyrosine hydroxylase immunocytochemistry. J Chem Neuroanat 18:117–133

Appeltants D, Ball GF, Balthazart J (2001) The distribution of tyrosine hydroxylase in the canary brain: demonstration of a specific and sexually dimorphic catecholaminergic innervation of the telencephalic song control nuclei. Cell Tissue Res 304:237–259.

Barker JM, Charlier TD, Ball GF, Balthazart J (2013) A new method for in vitro detection of bromodeoxyuridine in serum: a proof of concept in a songbird species, the canary. PLoS ONE 8(5): e63692. doi: 10.1371/journal.pone.0063692

Burek MJ, Nordeen KW, Nordeen EJ (1994) Ontogeny of sex differences among newly generated neurons of the juvenile avian brain. Devel Brain Res 78:57–64.

Colquitt BM, Merullo DP, Konopka G, Roberts TF, & Brainard MS (2021) Cellular transcriptomics reveals evolutionary identities of songbird vocal circuits. Science, 371, eabd9704

Daou A, Ross MT, Johnson F, Hyson RL, & Bertram R (2013). Electrophysiological characterization and computational models of HVC neurons in the zebra finch. J Neurophys 110(5):1227–1245.

DeWulf V, Bottjer SW (2005) Neurogenesis within the juvenile zebra finch telencephalic ventricular zone: a map of proliferative activity. J Comp Neurol 481:70–83.

Dutar P, Vu HM, & Perkel DJ (1998) Multiple cell types distinguished by physiological, pharmacological, and anatomic properties in nucleus HVc of the adult zebra finch. J Neurophys 80(4):1828–1838.

Fernstrom JD, Fernstrom MH (2007) Tyrosine, phenylalanine, and catecholamine synthesis and function in the brain. J Nutr 137:1539S–1547S.

Fortune ES, Margoliash D (1995) Parallel pathways and convergence onto HVc and adjacent neostriatum of adult zebra finches (Taeniopygia guttata). J Comp Neurol 360:413–441.

Haakenson CM, Balthazart J, & Ball GF (2020) Effects of inactivation of the periaqueductal gray on song production in testosterone-treated male canaries (Serinus canaria). 7(4) ENEURO.0048-20.2020 1–9

Hara M, Fukui R, Hieda E, Kuroiwa M, Bateup HS, Kano T, Greengard P, Nishi A (2010) Role of adrenoreceptors in the regulation of DA/DARPP-32 signaling in neostriatal neurons. J Neurochem 113(4):1046–1059.

Hemmings Jr HC, Walaas SI, Ouimet CC, Greengard P (1987) Dopaminergic regulation of protein phosphorylation in the striatum: DARPP-32. Trends Neurosci 10:377–383.

Jarvis ED, Scharff C, Grossman MR, Ramos JA, Nottebohm F (1998) For whom the bird sings: context dependent gene expression. Neuron 21:775–788

Keilani S, Chandwani S, Dolios G, Bogush A, Beck H, Hatzopoulos AK, Rao GN, Thomas EA, Wang R, Ehrlich ME (2012) Egr-1 induces DARPP-32 expression in striatal medium spiny neurons via a conserved intragenic element. J Neurosci. 32(20):6808–6818.

Kirn JR, & DeVoogd TJ (1989) Genesis and death of vocal control neurons during sexual differentiation in the zebra finch. J Neurosci 9(9):3176–3187.

Kirn JR, Alvarez-Buylla A, Nottebohm F (1991) Production and survival of projection neurons in the forebrain center of adult male canaries. J Neurosci 11(6):1756–1762.

Kirn JR, Fishman Y, Sasportas K, Alvarez-Buylla A, Nottebohm F (1999) Fate of new neurons in adult canary high vocal center during the first 30 days after their formation. J Comp Neurol 411:487–494.

Kosubek-Langer J, Schulze L, Scharff C (2017) Maturation, behavioral activation, and connectivity of adult-born medium spiny neurons in a striatal song nucleus. Front Neurosci 11:323. doi: 10.3389/fnins.2017.00323

Kubikova L, Wada K, Jarvis ED (2009) DA receptors in a songbird brain. J Comp Neurol 518:741–769.

Kubota M & Taniguchi I (1998). Electrophysiological characteristics of classes of neuron in the HVC of the zebra finch. J Neurophys 110(5):914–923.

Long MA & Fee MS (2008) Using temperature to analyze temporal dynamics in the songbird motor pathway. Nature, 456(7219):189–194.

Mooney R & Prather JF (2005) The HVC microcircuit: the synaptic basic for interactions between song motor and vocal plasticity pathways. J Neurosci, 25(8):1952–1964.

Nixdorf BE, Davis SS, & DeVoogd TJ (1989). Morphology of golgi-impregnated neurons in hyperstriatum ventralis, pars caudalis in adult male and female canaries. J Comp Neurol 284:337–349.

Nixdorf-Bergweiler BE, Bischof HJ (2007) A stereotaxic atlas of the brain of the zebra finch, Taeniopygia guttata: with special emphasis on telencephalic visual and song system nuclei in transverse and sagittal sections. Bethesda, MD: NCBI.

Ouimet CC, Greengard P (1990) Distribution of DARPP-32 in the basal ganglia: an electron microscopic study. J Neurocytol 19:39–52.

Pytte CL (2016) Adult neurogenesis in the songbird: region-specific contributions of new neurons to behavioral plasticity and stability. Brain Behav Evol 77:191–204.

Riters L, Eens M, Pinxten R, Ball GF (2002) Seasonal changes in the densities of a2-noradrenergic receptors are inversely related to changes in testosterone and the volumes of song control nuclei in male European starlings. J Comp Neurol 444-63-74.

Roberts TF, Hisey E, Tanaka M, Kearney M, Chattree G, Yang C, Shah NM, Mooney R (2017) Identification of a motor to auditory pathway important for vocal learning. Nat Neurosci 20(7):978–986.

Rochefort C, He X, Scotto-Lomassese S, Scharff C (2007) Recruitment of FoxP2-expressing neurons to Area X varies during song development. Dev Neurobiol 67(6):809–817

Ross MT, Flores D, Bertram R, Johnson F, & Hyson RL (2017). Neuronal intrinsic physiology changes during development of a learned behavior. eNeuro 25 September 2017, 4 (5) ENEURO.0297-17.2017

Ross MT, Flores D, Bertram R, Johnson F, Wu W, & Hyson RL (2019) Experience-dependent intrinsic plasticity during auditory learning. J Neurosci 39(7):1206–1221.

Scharff C, Kirn JR, Grossman M, Macklis JD, Nottebohm F (2000) Targeted neuronal death affects neuronal replacement and vocal behavior in adult songbirds. Neuron 25:481–492.

Scott BB, Lois C (2007) Developmental origin and identity of song system neurons born during vocal learning in songbirds. J Comp Neurol 502:202–214.

Scotto-Lomassese S, Rochefort C, Nshdejan A, Scarff C (2007) HVC interneurons are not renewed in adult male zebra finches. Eur J Neurosci 25(6):1663–1668.

Shea SD, Koch H, Baleckaitis D, Ramirez JM, & Margoliash D (2010) Neuron-specific cholinergic modulation of a forebrain song control nucleus. J Neurophys 103(2):733–745

Shaughnessy DW, Hyson RL, Bertram R, Wu W, Johnson F (2019) Female zebra finches do not sing yet share neural pathways necessary for singing in males. J Comp Neurol 527(4):843–855.

Shevchouk OT, Ball GF, Cornil CA, Balthazart J (2017) Studies of HVC plasticity in adult canaries reveal social effects of sex differences as well as limitation of multiple markers available to assess adult neurogenesis. PLoS One 12(1): e0170938. doi:10.1371/journal.pone.0170938

Singh UA & Iyengar S (2019). The expression of DARPP-32 in adult male zebra finches (Taenopygia guttata). Brain Struct Funct 224(8):2939–2972.

Sohrabji F, Nordeen EJ, Nordeen KW (1993) Characterization of neurons born and incorporated into a vocal control nucleus during avian song learning. Brain Res 620:335–338.

Sossinka R & Bohner J (1980) Song types in the zebra finch (*Poephila guttata castanotis*). Z. Tierpsychol. 53:123–132.

Svenningsson P, Nishi A, Fisone G, Girault JA, Nairn A, & Greengard P (2004). DARPP-32: an integrator of neurotransmission. Annu. Rev. Pharmacol. Toxicol. 44:269–296.

Tanaka M, Sun F, Li Y, Mooney R (2018) A mesocortical DA circuit enables the cultural transmission of vocal behavior. Nature 563(7729):117–120.

Tokarev K, Boender AJ, Claβen GAE, Scharff C (2016) Young, active and well-connected: adult-born neurons in the zebra finch are activated during singing. Brain Struct Funct 221:1833–1843.

Walaas SI, Greengard P (1984) DARPP-32, a DA- and adenosine 3’:5’-monophosphate regulated phosphoprotein enriched in DA-innervated brain regions. Regional and cellular distribution in the rat brain. J Neurosci 4(1):84–98.

Walton C, Pariser E, Nottebohm F (2012) The zebra finch paradox: song is little changed, but number of new neurons doubles. J Neurosci 32(3):761–774.

Wang N, Hurley P, Pytte C, Kirn JR (2002) Vocal control neuron incorporation decreases with age in the adult zebra finch. J Neurosci 22(24):10864–10870.

Wild JM, Williams MN, Howie GJ, Mooney R (2005) Calcium-binding proteins define interneurons in HVC of the zebra finch (Taeniopygia guttata). J Comp Neurol 483:76–90.

Wood WE, Roseberry TK, & Perkel DJ (2013). HTR2 receptors in a songbird premotor cortical-like area modulate spectral characteristics of zebra finch song. J Neurosci 33(7):2908– 2915.

Yao F, Zhang E, Gao Z, Ji H, Marmouri M, Xia X (2018) Did you choose appropriate tracer for retrograde tracing of retinal ganglion cells? The differences between cholera toxin subunit B and Fluorogold. PLoS ONE 13(10): e0205133. https://doi.org/10.1371/journal.pone.0205133

